# Kinematic analysis deconstructs the evolved loss of schooling behavior in cavefish

**DOI:** 10.1101/2020.01.31.929323

**Authors:** Adam Patch, Alexandra Paz, Karla Holt, Erik Duboue, Johanna E. Kowalko, Alex C. Keene, Yaouen Fily

**Affiliations:** Wilkes Honors College, Florida Atlantic University, Jupiter, FL 33458; Jupiter Life Science Initiative, Florida Atlantic University, Jupiter, FL 33458; Department of Biological Sciences, Florida Atlantic University, Jupiter, FL 33458

## Abstract

Fish display a remarkable diversity of social behaviors, from highly social to largely solitary. While social behaviors are likely critical for survival, surprisingly little is known about how they evolve in response to changing environmental pressures. With its highly social surface form and multiple populations of a largely asocial, blind, cave-dwelling form, the Mexican tetra, *Astyanax mexicanus*, provides a powerful model to study the evolution of social behavior. Given numerous morphological and behavioral differences between the surface and cave forms, a primary impediment to understanding how this behavior evolved is a lack of computational and statistical approaches that can precisely identify independent aspects of social behavior. Here, we use motion tracking and kinematic analysis to quantify social swimming patterns and argue that the absence of schooling in cavefish is not merely a consequence of their inability to see but rather a genuine behavioral adaptation that impacts the way they explore their cave environment. Surface fish school, maintaining both close proximity and alignment with each other. In the dark, surface fish no longer school, but we show that they still attempt to align and maintain proximity when they find themselves near another fish. Conversely, cavefish exhibit little preference for proximity or alignment, instead actively avoiding each other. Cavefish also slow down when more fish are present in the tank, which neither surface fish in the light or in the dark do. Using data-driven computer simulations, we show that those two traits – active avoidance and collective slowdown – are sufficient to shift the exploration strategy of cavefish from mostly-following-the-walls to exploring-the-entire-tank. Finally, we find that those differences in collective swimming patterns are largely consistent across independently-evolved cave populations, revealing an evolutionary convergence on this emergent social behavior.

**Author Summary:** The Mexican tetra fish offers a fascinating window into the evolution of schooling behavior. Its ancestral-like surface form is found in the rivers and lakes of Mexico and Texas and is highly social. Over the last million years, *A. mexicanus* repeatedly evolved a second, largely asocial cave form after colonizing a series of dark, underground caves. Here we use motion tracking technology to compare the collective displacement patterns of four populations and present evidence that the loss of schooling in cave populations (i) is a genuine example of parallel behavioral evolution rather than a mere consequence of not being able to see in the dark and (ii) could constitute a cave-specific exploration strategy.

## 1 Introduction

Social behaviors differ dramatically across species as a result of their unique evolutionary histories and ecological factors that include predation, foraging strategy, sexual selection, and available sensory inputs [27, 20, 13, 28]. Comparisons between closely-related species or populations within the same species are an essential source of insight into the variability of social behavior, its dependence on ecological factors, and its genetic and neural bases [14, 31]. Recent advances in motion tracking methods can dramatically refine such comparisons by providing highly-detailed, quantitative behavioral phenotypes [37, 34].

Fish schooling has been a particularly fertile ground for the application of motion tracking methods. In threespine sticklebacks, they were used to show that a marine population schools (fish maintain both proximity and alignment) whereas a benthic one shoals (maintains proximity only) and to identify some of the genes underlying this difference [44, 19, 18, 17]. In zebrafish, they allowed for quantification and categorization of the effects of 90 different genetic mutations on various aspects of schooling behavior, from the size of the school to its polarization [42]. When combined with physical theories of collective motion, motion tracking can also be used to infer mechanistic schooling models [16, 15, 29, 22, 4, 33]. This approach offers the fascinating prospect of simulating hypothetical mutations affecting the values of the model’s parameters and exploring their effect on the fish’s collective behavior [5] and their evolutionary fitness [21].

However, these advances have rarely been applied to those species most relevant to behavioral evolution. One such organism is the Mexican tetra, *Astyanax mexicanus*. It exists in two forms: a highly-social surface form, found in the rivers and lakes of Mexico and Southern Texas, and a much-less-social cave form found in at least 29 caves in the Sierra de El Abra and Sierra de Guatemala regions of Northeast Mexico. Surface and cave *A. mexicanus* populations inhabit very different environments, exhibit very different social behaviors, and can be interbred to identify the genetic underpinnings of their behavioral differences. Additionally, the cave form of *A. mexicanus* has evolved multiple times independently [23]. Finally, the loss of schooling is just one in a series of morphological and behavioral differences between the surface and cave forms including eye loss, increased sensitivity of the mechanoreceptive lateral line, and a reduction in aggression [46, 43, 26, 11, 32], all of which may directly or indirectly disrupt schooling behavior and, in turn, encourage new social interactions. Vision-deprived surface fish do not school, suggesting the lack of visual inputs may explain the lack of schooling in cavefish. However, some surface-cave hybrids that respond to visual cues do not school either, suggesting the loss of schooling may also have a vision-independent component [32]. Unfortunately, the relatively simple approach used to quantify schooling in previous studies [32] makes it difficult to compare the behaviors of cavefish and vision-deprived surface fish beyond their common lack of schooling. Thus, *A. mexicanus* provides both the opportunity to explore some fundamental-yet-understudied aspects of the evolution of schooling behavior and a system in which motion tracking methods can help collect previously unavailable information.

In this paper, we use a motion-tracking-based schooling assay to address four questions related to schooling and its evolution in *A. mexicanus*: (i) What is an appropriate set of metrics to quantify social swimming in *A. mexicanus* beyond schooling/not schooling? (ii) Can those metrics clarify the connection between the loss of schooling and the loss of vision? (iii) Can those metrics identify differences between different cave populations? (iv) What fundamental behavioral components should a mechanistic model of *A. mexicanus* swimming include?

## 2 Results

### 2.1 Boundary effects and the importance of tank design

Although surface and cave populations of *A. mexicanus* have been studied extensively for evolved differences in locomotion, schooling, and sleep, those assays typically involved contrived environments that may not reflect motion under natural conditions including fish interacting through a transparent partition or swimming with a school of artificial fish and small, often rectangular, tanks that bias the fish’s orientation and turning behavior [47, 32, 35, 45, 30, 32, 44]. Clean turning statistics are particularly important for mechanistic schooling models, which normally start by modeling unprompted changes of speed or orientation, i.e., changes that are not triggered by the proximity of a wall or other fish, before modeling the effect of the walls (by comparing trajectories near/far from the wall) and the other fish (by comparing trajectories near/far from other fish). With this in mind, we first examine the swimming dynamics of single fish.

Individual fish from surface, Pachón, Tinaja, or Molino populations were filmed in a circular tank with a diameter of 111 cm (about 20 fish body lengths) (Fig 1a and b). The circular shape was picked because it does not favor any overall orientation. The tank was filled to a depth of 10 cm so that motion was mostly two-dimensional and could be filmed from above with minimal information loss. Fish were filmed for 20 minutes following a 10-minute acclimation period.

**Figure 1:**
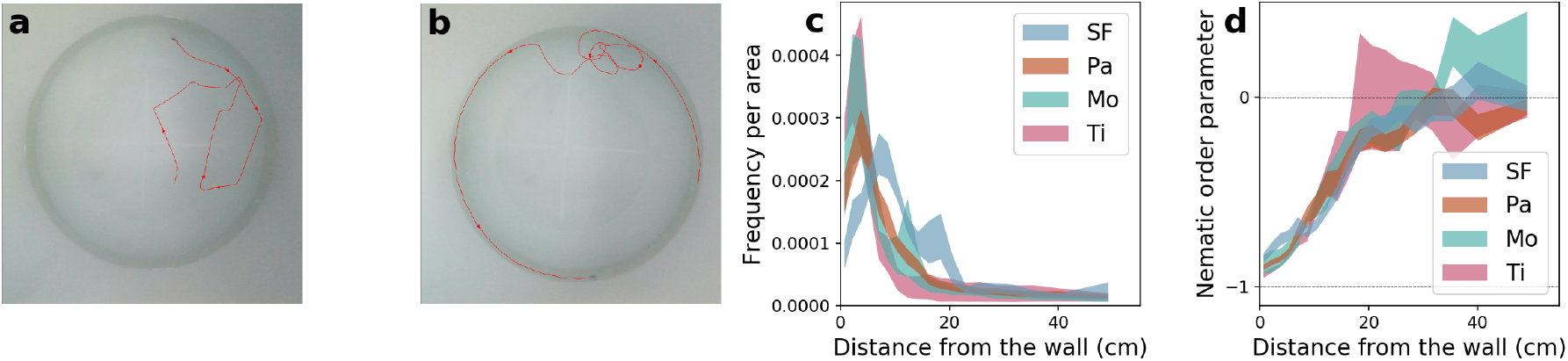
(a) Snapshot from a video of a single surface fish. The red line is the trajectory over the 25 seconds leading to the snapshot. (b) Similar snapshot and 25 s-trajectory from a video of a single Pachón cavefish. (c) Fish density probability per unit area as a function of the distance from the wall for single fish trials for each population. The distance varies between 0 cm (right against the wall) and 55.5 cm (center of the tank). (d) Average nematic alignment parameter relative to the wall (cos(2*θ*) where *θ* is the angle between the fish’s heading and the normal to the closest wall) as a function of the distance to the wall. Values near −1 close to the wall indicate strong alignment parallel to the wall. Values near 0 in the center suggest no favored orientation. The width of the curves in c and d corresponds to one standard error on either side of the mean. This convention is used for probability distributions throughout the paper.

To assess how far the effect of the walls influences the dynamic movements of the fish, we measured the statistics of their distance to and orientation with respect to the nearest wall (Figure 1cd). All populations spend the vast majority of their time near the wall (Figure 1c, all densities show a pronounced peak near the wall). However, cavefish stay closer to the wall than surface fish: cavefish peak sharply between 2–5 cm, while surface fish have a broader peak at 7 cm. For every population, the density per unit area reaches a plateau starting between 10 and 20 cm from the wall, meaning the density no longer depends on the distance to the wall, i.e., the tank is large enough to observe a bulk density. Further, while fish near the wall display nematic wall-alignment parameter values near *−*1, demonstrating that they stay mostly parallel to the wall, fish located more than 20 cm from the wall exhibit values near 0, suggesting they orient independently of the wall (Figure 1d).

Overall, this shows that the effect of the walls remains strong up to at least 20 cm from the wall. When designing a tank to quantify the dynamics of surface and cave *A. mexicanus* in situations where they are not dominated by the wall, one should therefore use a tank larger than 40 cm in diameter. Additionally, some results from the literature may have to be reinterpreted. For example, Sharma et al. found that surface fish occupy the entirety of a 30 cm-diameter tank [38] whereas cavefish stay near the wall. What our results suggest is that surface fish do prefer to stay near the wall, however the entirety of a 30 cm-diameter tank is within 15 cm of the wall, which according to Figure 1c is still near the wall to a surface fish.

### 2.2 Group size affects speed regulation and the nature of turns

Schooling and shoaling are defined by the way the fish in a group position and orient themselves with respect to each other. However, the presence or absence of other fish can also affect a fish’s overall swimming behavior, whether or not it is near another fish. For example, surface fish exhibit a stress response to isolation which makes them seek the bottom of their tank [6]. Although depth is not significant in our shallow tank, changes in the mean speed or turning statistics are. Cataloging such effects is also important to guide future mechanistic models. Thus, we calculated the effects of group size on the statistics of a fish’s speed and turning speed.

#### Isolated surface fish pause

The most striking effect of group size is that surface fish in groups of 1, 2, and 5 occasionally stop moving, as evidenced by the sharp zero-speed peak in their speed distributions (Figure 2a). The area under the peak represents the average fraction of the time the fish were inactive. The base of the peak is consistently located around 1 cm/s, providing a natural definition of active (moving faster than 1 cm/s) versus inactive (slower than 1 cm/s) fish. The fraction of the time surface fish were active shows a clear trend, from 73% in single-fish trials up to close to 100% in groups of 10 (Figure 2b). In contrast, cavefish keep moving regardless of the number of fish in the tank (Figure 2b).

**Figure 2:**
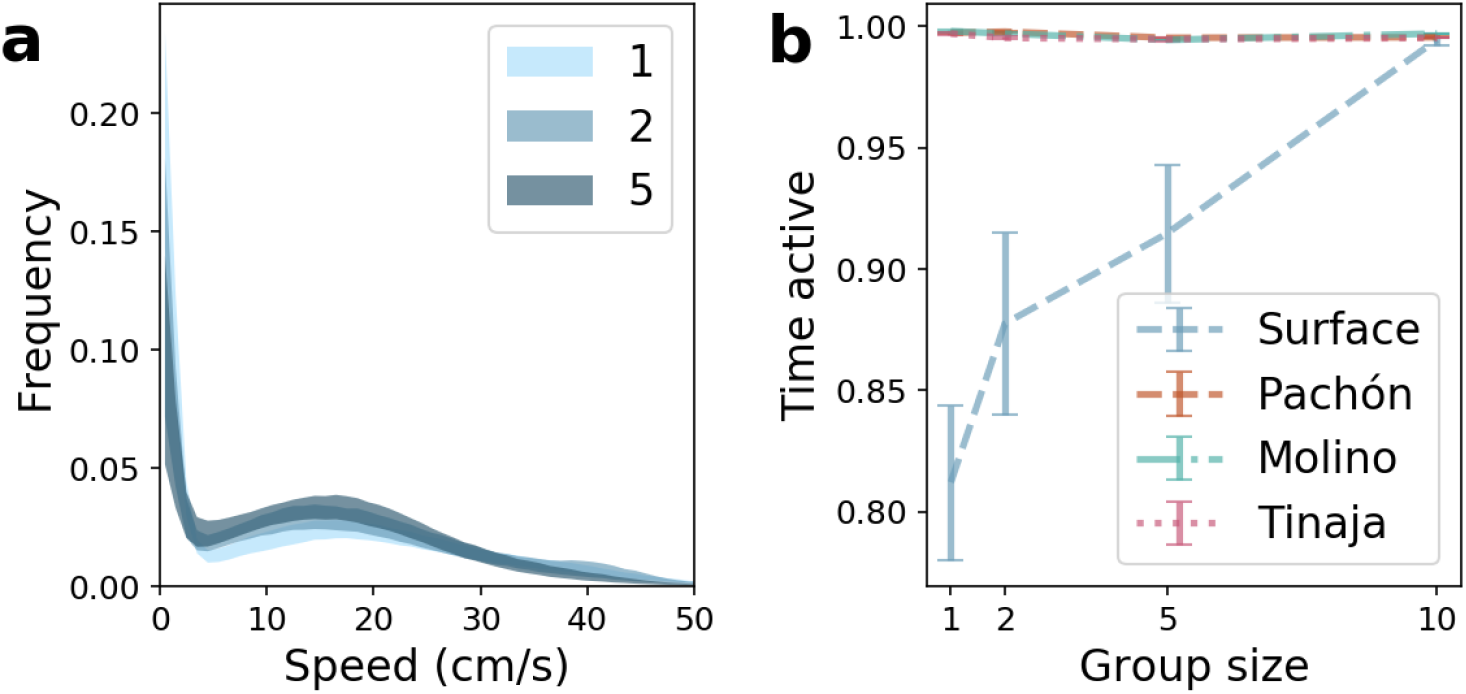
(a) Speed distribution of surface fish as a function of group size. A sharp peak near zero indicates fish are not moving. The area under the peak is the fraction of the time the fish are not moving. (b) Fraction of time spent moving faster than 1 cm/s (“active”) as a function of group size. Cavefish are almost always moving, whereas surface fish are less active when they are more isolated.

#### Inactivity cut

Throughout the rest of the paper we focus on swimming properties in the active state, i.e., we exclude the inactive data (speed<1 cm/s) from all subsequent distributions and averages. In practice this *inactivity cut* only affects groups of 1, 2, and 5 surface fish, i.e., those shown in Figure 2a.

#### Cavefish slow down in larger groups

Once inactive frames have been removed to focus on the speed measured while swimming, we see that the speed distribution of surface fish depends very little on the number of fish in the tank (Figure 3a). Conversely, cavefish noticeably slow down as the number of other fish increases (Figure 3b). Perhaps surprisingly, this slowdown is not limited to encounters with other fish but rather reflects the overall behavior of the fish both during and between encounters, as evidenced by the speed distribution shifting as a whole rather than developing a second, slower peak (Figure 3b). All three cave populations slow down in a similar way with increased group size (Figure 3c).

**Figure 3:**
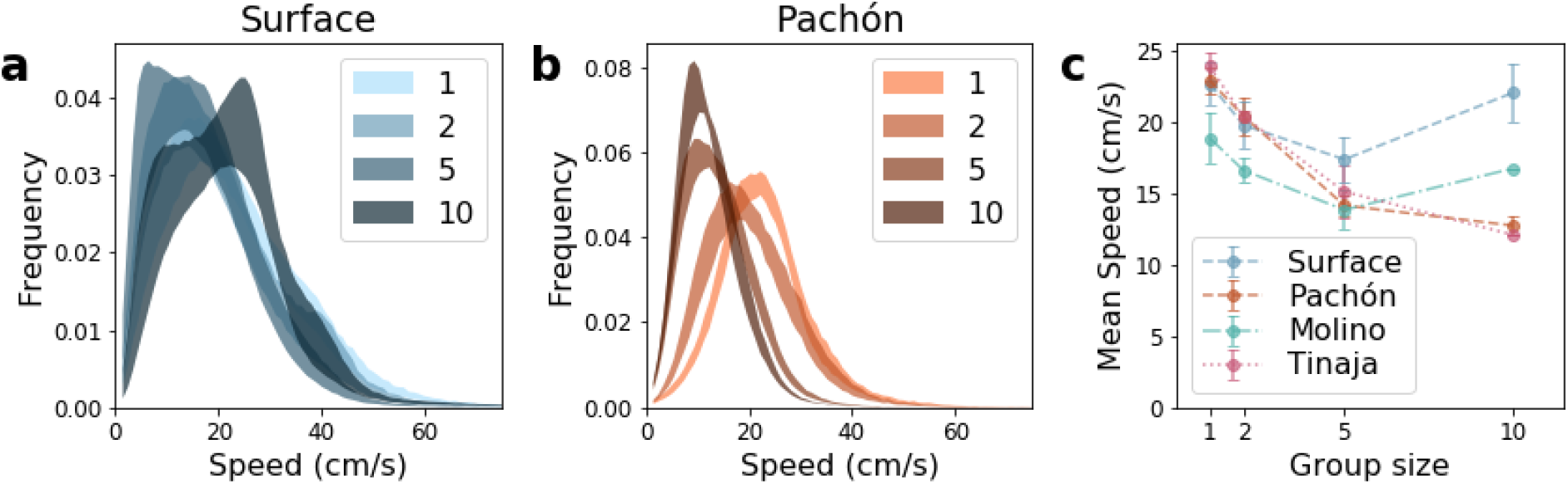
Speed distribution as a function of group size for actively swimming (speed>1 cm/s) (a) surface fish and (b) Pachón cave populations. (c) Mean speed as a function of group size for all types shows a convergent social behavior in all cave populations as cavefish swimming speed decreases with group size while surface fish maintain a constant speed regardless of group size. Tinaja and Molino speed distributions resemble those of Pachón and are shown in Figure S1.

#### Surface fish turn more abruptly in smaller groups, cavefish in larger groups

With the speed statistics, the other fundamental quantity needed to characterize the undisturbed (by walls or other fish) motion of a fish is its distribution of turning speed (angular speed). Mechanistic models often use one of two assumptions: that the angular speed distribution follows a normal distribution, or that there is an alternation of straight bouts and instantaneous changes of direction. The former has been used to model barred flagtails [16]. The latter has been observed in rummy-nose tetra [4] and larval zebrafish [9].

Isolated surface fish spend a significant fraction of their time going straight (zero-angular-speed peak in Figure 4a), and much of their turning is sudden (plateau and slow decay in Figure 4b),consistent with the sharp turns in the sample trajectory of Figure 1a. In contrast, isolated cavefish turn more often (smaller zero-angular-speed peak in Figure 4c) and more gradually (peak around 4 rad/s then quick decay in Figure 4d), consistent with the sample trajectory shown in Figure 1b. When the number of fish in the tank increases, the turning statistics of surface and cave fish change in opposite ways. In surface fish, increasing group size decreases both straight motion and quick turns in favor of gradual turning. In Pachón, larger groups exhibit less gradual turning, more straight motion, and more quick turning. Interestingly, most of the change we see in Pachón fish occurs between 1 and 2 fish, suggesting Pachón fish register the presence of another fish and modify their behavior even if their encounters with the other fish are somewhat infrequent.

**Figure 4:**
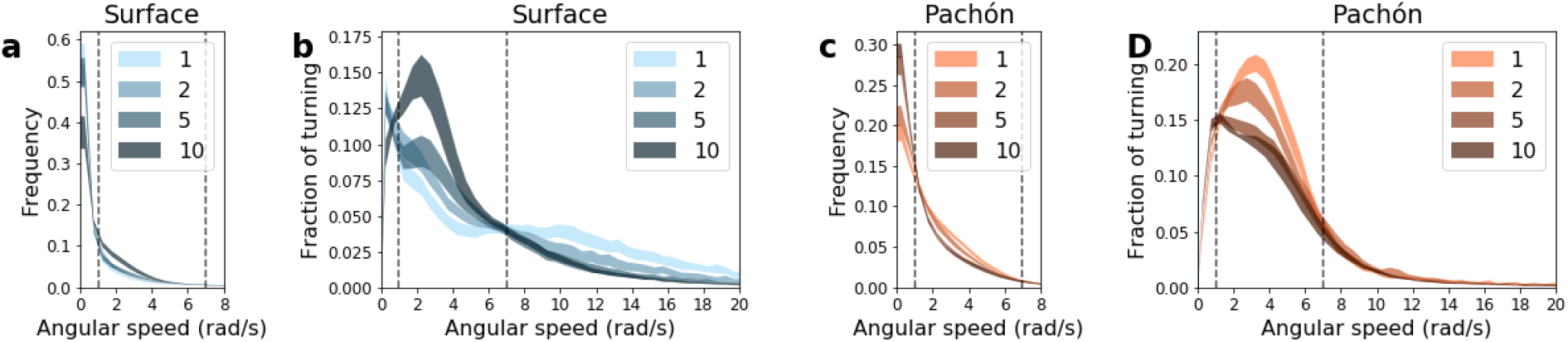
Turning distributions as a function of group size for (a,b) surface fish and (c,d) Pachon. (a) and (c) show the relative time spent turning at each speed. (b) and (d) show the relative fraction of the total angle turned that was turned at each speed. Although they contain the same information, the former better highlights the proportion of straight bouts whereas the latter better highlights which type of turn (quick or slow) contributes to the most turning (the largest angle turned). The role of group size is best understood in terms of three broadly defined ranges of turning speed: mostly straight (<1 rad/s; includes following the wall), gradual turning (1–7 rad/s), and sudden turning (>7 rad/s). The edges of those ranges are shown as vertical dashed lines in all four panels. Distributions for Tinaja and Molino are shown in Figure S2.

In terms of mechanistic modeling, perhaps the most significant finding here is that surface fish shift from an instantaneous-turn type of dynamics when isolated to a more continuous type of turn in larger groups. These two behaviors are usually described with different types of mathematical models. As a result, treating the two-fish dynamics as two single-fish dynamics plus interaction forces and torques may not be sufficient to capture the transition from isolated swimming to group swimming.

### 2.3 Surface fish school, vision-deprived surface fish try, cavefish do not

When presented with a moving school of artificial fish, surface fish tend to follow it whereas cavefish and vision-deprived surface fish do not [32]. However, little is known about the structure of surface fish schools or the spatial correlations in non-schooling groups of cavefish or vision-deprived surface fish. If the lack of schooling in cavefish is mainly caused by their lack of vision, we expect groups of cavefish and groups of vision-deprived surface fish to exhibit similar spatial arrangements. To address those questions, we consider the statistics of the distance between fish and the angle between their headings (Figure 5).

**Figure 5:**
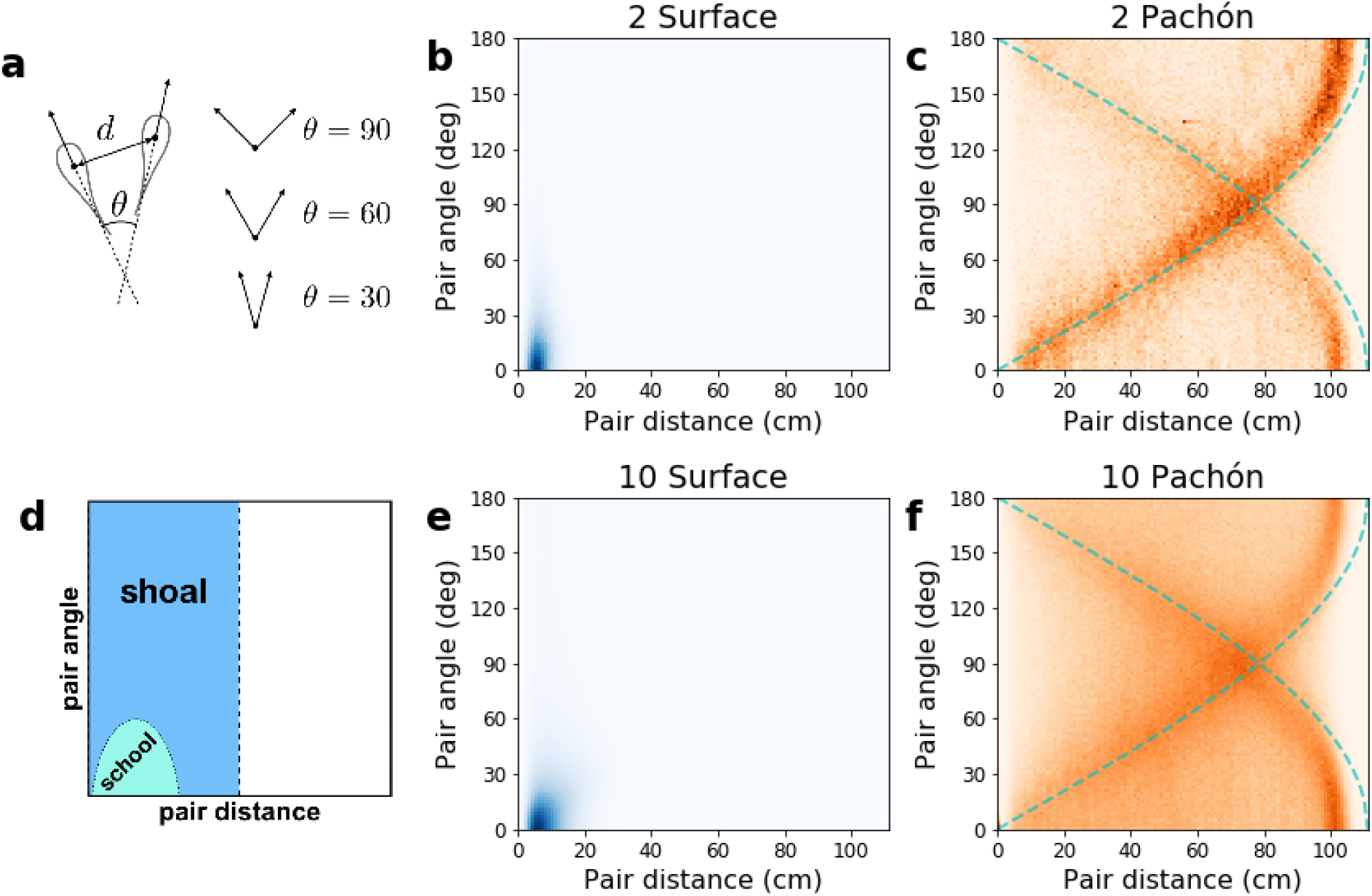
(a) Definition of the pair distance *d* and pair angle *θ* between two fish. (b, c, e, f) Joint probability density of pair distance and pair angle for all pairs of fish. (d) Qualitative expectations in common cases. For a school, short distances and small relative angles yield a peak in the bottom left corner. For a shoal, short distances and a wide range of angles yield a vertical band on the left side. For a group that does neither, we expect the distribution to spread over the entire available range of distances and angles. (b, e) Groups of surface fish exhibit close proximity and tight angles indicative of strong schooling. (c, f) Group of Pachón cavefish exhibit neither schooling nor shoaling. The cyan dashed lines show the expectation for fish following the wall (not necessarily the same part of it). Groups of 10 do not follow the walls as much. Comparable probability densities shown for all group sizes and populations (including Tinaja and Molino) in Figures S3, S4, and S5.

#### Surface fish form tight schools

For surface fish, we find a strong preference for both proximity and alignment, indicating genuine schooling rather than mere shoaling (Figure 5be, the joint probability distribution of inter-fish distance and angle peaks in the bottom left corner corresponding to short distance and small angle). In groups of two, the distance between the fish is less than 15 cm (3 average body lengths) 96.1% of the time and the angle between their headings is less than 45° 85.5% of the time. In groups of ten, the distance between any two fish is less than 15 cm 75.5% of the time and the angle between their headings is less than 45° 81.5% of the time.

#### Cavefish do not school or shoal

Unlike surface fish, the joint probability distribution of inter-fish distance and angle in cavefish (Figure 5cf) reveals neither a preference for proximity nor one for alignment. The most striking pattern it exhibits is a pair of arches due to the fish’s tendency to follow the walls (the cyan dashed lines represent two points moving along the surface of the wall at all times). Interestingly, this pattern subsides in larger groups, suggesting the fish do not swim along the walls as much.

#### Darkness disrupts schooling, but surface fish still align at short range

To find out whether the loss of schooling in cavefish can be explained by their lack of vision, we measure the speed distribution and the joint probability of inter-fish distance and angle in surface fish in the dark (Figure 6). Surface fish in the dark kept moving over the entire trial, similar to cavefish, but unlike surface fish in the light, which occasionally stop moving when in smaller groups (up to 5 fish, Figure 2). Conversely, the mean swimming speed of surface fish in the dark is mostly independent of the number of fish, similar to surface fish in the light but unlike cavefish, which swim slower in larger groups (Figure 6a). Finally, the probability distribution of inter-fish distance and angle of surface fish in the dark exhibits both schooling and non-schooling features (Figure 6b). Similar to the cavefish distribution (Figure 5f), it spreads over the entire available range of distances and angles, indicating there is no long-lived, compact school like the ones we see in the light. On the other hand, the distribution does exhibit a peak at small distance and small angle (bottom left corner of Figure 6b and left side of Figure 6c) showing that vision-deprived surface fish still have a preference for proximity and alignment, even though those are no longer sufficient to form a school. Consistent with this interpretation, direct inspection of the video data shows pairs of surface fish in the dark swimming together for a brief period of time but drifting apart before more fish can join this “proto-school”.

**Figure 6:**
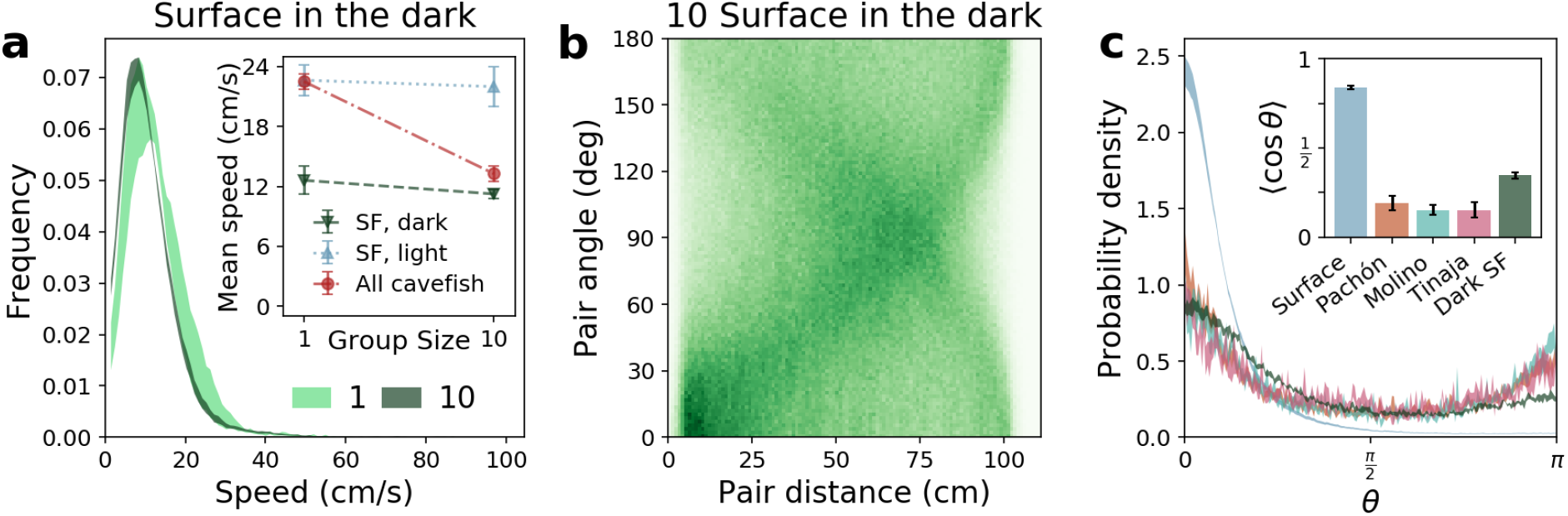
Summary of dark trials for surface fish, alone and in groups of ten. (a) The speed distribution shows slower swimming than in the light, but little change as a function of group size. The probability density of relative distance and angle exhibits both a spread over the entire tank and a low peak at small distance and angle, indicating locally coordinated swimming but no fully formed school. (c) The probability density of the pair angle *θ* when the two fish are near each other (within 10 cm) confirms short-range alignment. Cavefish distributions are roughly symmetric around *θ* = 90° indicating no preference for alignment vs anti-alignment. Conversely, surface fish distributions show a preference for acute angles (*θ* < 90°). The inset shows the average of cos *θ*, which measures whether the two fish tend to move in the same direction (⟨cos *θ*⟩ > 0) or in opposite directions (⟨cos *θ*⟩ < 0), showing that cavefish are significantly less likely to be aligned than surface fish in the dark.

In mathematical models of collective motion, inter-individual interactions favoring proximity and alignment compete with noise (random, unprompted course changes). If the interactions are strong enough, global order emerges, i.e., a well-ordered school. When the interactions are weaker than the noise, one still expects local order, i.e., an increased probability for nearby fish to align and/or stay together, which is precisely what we observe in surface fish in the dark. What is more, decreasing the range of the interaction can be sufficient to prevent system-wide alignment [7]. Therefore, our interpretation of the surface-fish-in-the-dark data is that the fish do attempt to school, however the loss of vision reduces their range of perception and prevents them from schooling effectively. This is in contrast to cavefish, which show almost no preference for alignment (Figure 6) and even actively avoid each other (Figure 7 and section 2.4 below).

**Figure 7:**
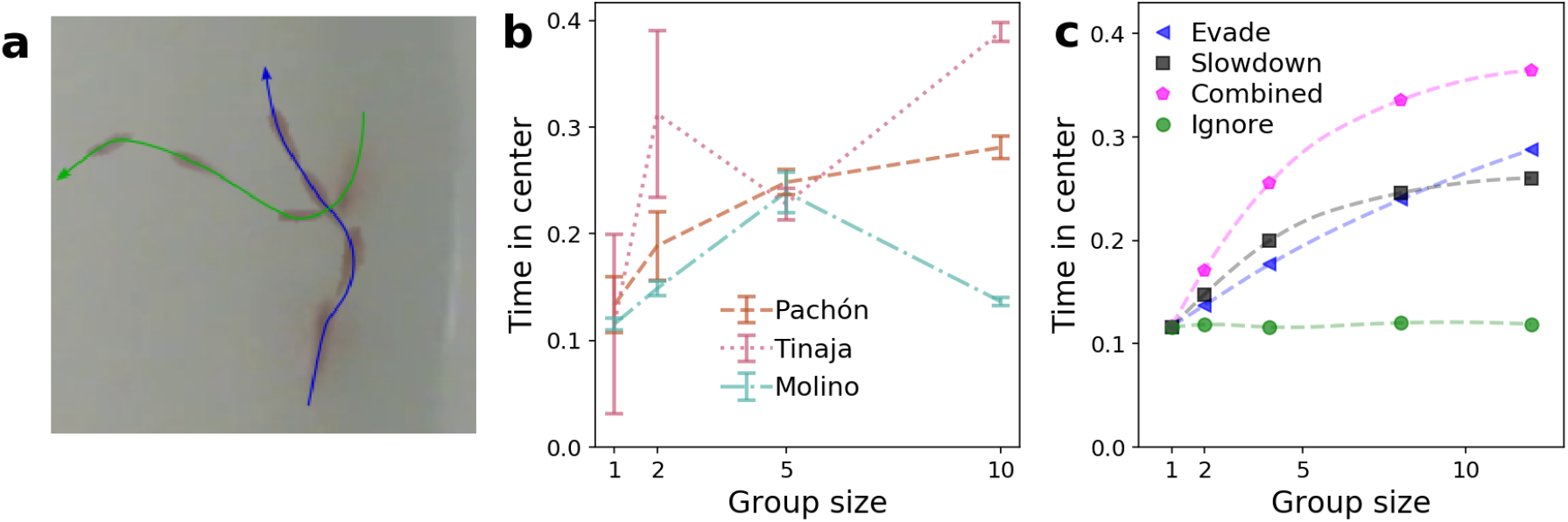
(a) Example of an encounter between two Pachón leading to a major change of direction. (b,c) Fraction of the time spent in the inner half of the tank (within 39.2 cm of its center, which represents half of its area) as a function of group size (b) in experiment and (c) in simulations. Persistent motion alone favors the outer half, but both swimming slower (which cavefish do in larger groups) and turning more (which evasive maneuvers contribute to) can weaken this effect. Our simulations suggest that when individuals ignore each other, they do not spend more time in the center of the tank. On the other hand, evasive interactions and slowing down with social density results in more time away from the walls.

### 2.4 Avoidance behaviors lead cavefish away from walls

Wall-following may serve an exploratory function in cave environments [38, 35]. If the walls get overcrowded, however, there may be a benefit to exploring away from them. The dimming of the wall-following arch pattern in Figure 5f compared to Figure 5c suggests cavefish do not follow the tank’s wall as much when more fish are present in the tank. To investigate this change, we measure the fraction of the time the fish spend in the center of the tank and use data-driven computer simulations to explore possible explanations. As expected from Figure 5cf, the fraction of the time cavefish spend in the inner half of the tank (thereafter *time in the center*) trends upwards with increased group size (Figure 7b).

Direct observation of cavefish motion suggests two possible mechanisms for this increased time away from the walls. The first comes from the observation that when cavefish get too close, they often react with an evasive maneuver. This tends to happen when they follow the wall in opposite directions, where a turn can send one or both off the wall and towards the center (Figure 7a). The second possible mechanism follows our observation that cavefish slow down with increased group size (Figures 3c and 6a-inset). Both increased turning and slower motion decrease the persistence length (the average distance traveled before turning), which controls the generic preference for the wall observed in many persistently moving systems including *E. coli* bacteria, sperm cells, and abstract self-propelled particle models even in the absence of explicit attraction to the wall [8, 39, 10].

To test those possible mechanisms, we simulated several variants of a simplified model of cavefish swimming (Section 4.8). The *evade* model simulates fish turning away from each other upon closeencounter. The *slowdown* model adjusts the swimming speed as a function of group size according to the mean speed observed in Pachón fish (Figure 3c). We combines these two in our *combined* model. The *ignore* model has neither evasion nor slowdown and plays the role of control model. All three models (*slowdown*, *evade*, and *combined*) exhibit an increase of the time in the center as the group size increases, and this effect is comparable to the one observed experimentally (Figure 7bc).

## 3 Discussion

In this paper, we systematically investigated the collective behavior of *A. mexicanus* individuals drawn from three populations of independently-evolved cavefish and compared them to their extant surface fish relatives, first in the light, then in the dark to simulate sight loss. To generate semi-realistic conditions and allow for the statistical analysis of the spatial structure of groups of fish, we let the fish interact freely in a circular tank that is much larger than those used in previous studies of locomotion and social behavior. We then applied automated tracking [40, 3] to quantify locomotion and interactions between individuals. We identified a series of collective behavioral differences between surface and cave populations that are common to the three cave populations, cannot be explained solely by their common lack of sight, and impact the way they navigate their environment, thus revealing evolutionary convergence on shared mechanisms of collective behavior.

The first point we clarified is the relationship between the loss of sight and the loss of schooling in cavefish. An earlier study of schooling in surface and cave fish suggested that loss of vision plays a large role in the loss of schooling in cavefish, but is probably not the only cause, with additional vision-independent mechanisms also at play [32]. A key piece of evidence for the existence of such vision-independent causes is the absence of schooling in some surface-cave hybrids with seemingly intact visual systems. Here we provide a new, independent piece of evidence that the lack of schooling in cavefish is a behavioral change that cannot be explained solely by the lack of visual input. By going beyond the schooling/non-schooling dichotomy and quantifying the dynamic structure of groups of *A. mexicanus*, we are able to show that the absence of schooling seen in cavefish and the one seen in vision-deprived surface fish are of very different natures: the latter show a preference for proximity and alignment with each other at close distances, suggesting they attempt to school but are unable to, whereas the former exhibit very little preference for proximity or alignment. It follows that the loss of schooling in cavefish is not just a matter of lacking the necessary visual input.

Our quantification of displacement patterns in *A. mexicanus* also reveals that cavefish actively avoid each other in at least two ways. First, cavefish swim slower when more fish are present in the tank, which mechanically decreases the frequency of their encounters with other fish. Importantly, this is not an instantaneous slowdown based on the current number of nearby fish (a behavior which actually promotes aggregation rather than avoidance [12]). Rather, cavefish seem to respond to the total number of fish in the tank. It is unclear whether that is achieved by some form of remote sensing (e.g. overall level of vibration in the tank), counting the number of recent encounters, or other means. Second, close encounters with other fish often lead to exaggerated turns whose amplitudes seems more consistent with evasive behavior than simple collision avoidance. Regardless of the mechanism(s), those avoidance behaviors support the idea that cavefish, unlike vision-deprived surface fish, make no attempt to school.

Interestingly, both avoidance mechanisms (slowdown and evasive turns) promote exploration away from the walls in cavefish. We therefore suggest that they constitute a form of collective response by which groups of cavefish adjust their exploration strategy to the crowdedness of their environment. Depending on the spatial distribution of resources in cave environments, this could provide a fitness advantage that may have contributed to the de-evolution of schooling in cavefish. Future ecological data on the shape of cave environments and the spatial distribution of cavefish-relevant resources in them would be of great interest and could be fed into our computer simulations to assess the potential fitness benefit of various exploration strategies.

Another goal of our fine quantification of collective swimming patterns in *A. mexicanus* was the identification of potential differences between cave populations. Although there are hints Molino fish may not avoid each other as much as Pachón or Tinaja fish, which we hope to confirm in a future larger study, the most striking outcome of this comparison is how remarkably similar the Pachón, Tinaja, and Molino cave populations behave. The three populations did not merely all stop schooling: none of them pause when alone, they all slow down when more fish are present in the tank, they all exhibit rounder turns, and none of them seem to seek proximity or alignment. In many cases, even probability distributions measured from different populations are within each other’s error bars.

The last goal of our study was to make steps towards a mechanistic model of collective swimming in *A. mexicanus*. On the one hand, our results suggest that *A. mexicanus* may be less amenable to highly accurate modeling than some other species, largely because the presence of other fish modifies many aspects of the swimming dynamics, from the speed distribution (including the propensity to swim at all) to the nature of the turns (including surface fish shifting from occasional sharp turns to frequent gradual ones). On the other hand, the mathematical modeling of schooling behaviors is a very active field of research, one in which recent advances may apply to *A. mexicanus* and one in which *A. mexicanus* could push new paradigms. Either way, our analysis of the role of the crowd-induced slowdown and evasive turns in cavefish on their exploration of the tank shows that even simple models that gloss over most of those subtleties can provide valuable insight.

Beyond those specific points, our work illustrates the benefits of generating highly quantitative behavioral phenotypes in evolutionarily tractable model organisms. The ability to quantify diverse aspects of locomotion behavior including velocity, wall proximity, angular velocity, and spatial relationships between fish represent an advance over previous analyses that quantified simple proximity to neighbors or exclusively examined locomotor activity [32, 47]. Our assay and analysis pipeline can be readily adapted to study other behaviors with a kinematic signature, e.g., aggression, sleep, stress and chemosensory behaviors [11, 1, 6, 25]. Fine quantification of those behaviors in surface-cave hybrids should yield new insight into their genetic basis. The recent development of a brain atlas and the mapping of the neuromorphology and neuronal activity in the same four populations discussed in this paper also offers interesting prospects, particularly if we can confirm behavioral differences between cave populations and connect them to morphological and neuronal differences [24]. Together, those combinations illustrate the potential of *A. mexicanus* as a model organism for the study of behavioral evolution as high-quality data sets become available all along the chain of events that leads from genetics, to morphology, to neuronal activity, to behavior.

## 4 Materials and Methods

### 4.1 Experimental Procedure

#### 4.1.1 Husbandry

Animal husbandry was carried out as previously described [41, 2]. All protocols were approved by the IACUC of Florida Atlantic University. Fish were housed in groups in Florida Atlantic University core facilities at a water temperature of 23 *±* 1 °C in 10 or 20 gallon tanks on a 14:10 hour light-dark cycle. Light intensity was maintained between 24 and 40 lux. Experimental fish were bred in the lab from descendants of either cavefish originally collected from the Pachón, Molino, and Tinaja caves or from two different surface populations originating from Texas or Mexico.

#### 4.1.2 Behavioral experiments

All experimental fish were adults housed in groups of 5-10 fish for 10 gallon tanks and 10-25 fish for 20 gallon tanks. Tanks and groups were chosen to avoid repeatedly assaying the same groups of fish and a minimum of two weeks was allowed before assaying fish from a previously used tank. Fish to be assayed were carried to a designated behavior room in a 2.5 L carrier tank and were then gently netted into the experimental arena and allowed to acclimate for 10 minutes.

Experiments were conducted in a round tank (111 cm diameter × 66 cm height) filled to a depth of 10 cm with water taken directly from the tank system. A video camera (Genius WideCam F100, Dongguan Gaoying Computer Products Co., Guangdong, China) was affixed to a custom-built PVC stand that allowed recording from above the center of the tank. Lighting was provided via four white 75-watt equivalent halogen light bulbs (Philips A19 Long Life Light Bulb, Amsterdam, Netherlands) mounted in clamp lights with 5.5 in shades (HDX, The Home Depot, Georgia, United States) to diffuse light. For experiments conducted in the dark, videos were collected by removing the infrared filter from within the camera, and whitelight bulbs were replaced with 940 nm Infrared bulbs (Spy Camera Specialists, Inc., New York, United States) and supplemented with four additional infrared lamps (IR30 WideAngle IR Illuminator, CMVision, Texas, United States). Videos were collected at 30 fps using OBS Studio (Open Broadcaster Software).

### 4.2 Data collection

Data was first collected in videos that begin recording when fish are released in the tank. The first 10 minutes of video are treated as acclimation time that is later discarded. Analysis focuses on the following 20 minutes. Table 1 summarizes the number of trials and the number of post-acclimation minutes collected for each group size and fish type. Trials were generally run for more than 30 minutes total. However, we have also included a small subset of videos (6 surface and 3 Pachón) despite only having 10 minutes post-acclimation.

**Table 1:** Summary of trials collected showing total trials collected and total time used in our analysis. Time is reported in minutes and represents video collected post-acclimation and prior to the end of the 20-minute time window used for analysis.

| Group Size | Surface (light) |  | Pachón |  | Tinaja |  | Molino |  | Surface (dark) |  |
| --- | --- | --- | --- | --- | --- | --- | --- | --- | --- | --- |
|  | trials | time | trials | time | trials | time | trials | time | trials | time |
| 1 | 20 | 363 | 20 | 382 | 3 | 60 | 3 | 60 | 3 | 60 |
| 2 | 10 | 182 | 9 | 162 | 3 | 60 | 3 | 60 | - | - |
| 5 | 12 | 228 | 10 | 192 | 3 | 60 | 3 | 60 | - | - |
| 10 | 10 | 197 | 4 | 80 | 1 | 20 | 1 | 20 | 4 | 80 |

### 4.3 Tracking

The positions and orientations of the fish were extracted from the videos using a custom python tracking library inspired by the tracktor library [40]. For each frame of each video, the library (i) identifies regions that are darker than their surroundings using adaptive brightness thresholding, (ii) discards regions whose area is not consistent with a fish, (iii) computes the center and orientation of each region, (iv) predicts where each of the fish found in the last frame will be in the current frame, (v) identifies the region corresponding to each fish by minimizing the distance between the centers of the regions and the predicted fish locations.

The tracking does not distinguish between the front and the back of the fish. That information is added after computing the velocity by assuming the fish always move forward.

### 4.4 Tank detection and pixel-centimeter conversion

Our custom tracking program includes an interactive tool to specify the circle corresponding to the edge of the tank. Fish coordinates are then converted to centimeters using the known physical diameter of the tank (111 cm).

### 4.5 Uncertainty due to swimming depth

Although we keep the water depth in the tank small (10 cm), fish can still move up or down by a few centimeters. Given the depth of the water, the height of our camera, and the assumption that fish are generally swimming at half depth, radial positions may be off by as much as 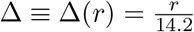. At the wall, this is as much as 3.87 cm, whereas in the center it is zero.

### 4.6 Kinematics

Time derivatives are computed using a standard second order central finite difference scheme. For example:

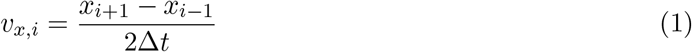

where *x*_*i*_ is the *x* coordinate of the position in frame *i*, *v*_*x,i*_ is the x coordinate of the velocity in frame *i*, and Δ*t* = 0.033 s is the duration of each frame (one over the frame rate).

To compute the angular velocity, the angle difference is first brought back into the (−*π, π*] interval:

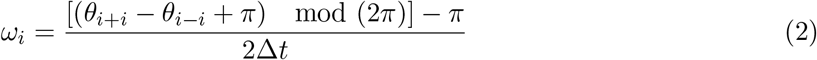

where *θ*_*i*_ is the angle the fish makes with the *x*-axis in frame *i*, *ω*_*i*_ is the angular speed in frame *i*, and mod is the modulo operator.

### 4.7 Data filtering

Tracking errors occasionally result in unrealistic speeds or angular speeds. To mitigate the effect of such errors on the overall analysis, we exclude frames in which a fish is moving faster than 100 cm/s or turning faster than 30 rad/s. The cut only affects the fish exhibiting the rapid motion, i.e., the frames are still used to analyze the other fish’s motion. We also exclude occlusion events, i.e., frames in which two fish overlap and cannot be distinguished from each other by the tracking software. For both cuts, we also excludes the three frames preceding the event and the three frames following it to avoid contaminating the kinematic quantities with data from the invalid interval. The inactivity cut discussed in section 2.2 also uses a three-frame buffer on either side of every inactive interval. Furthermore, if more than half of the frames in a trial are inactive, we remove the entire trial, in accordance with Ref. [32].

### 4.8 Simulations

#### 4.8.1 Models

Here we present a minimal model evasive active Brownian particles (ABPs) as preliminary consideration of collective effects that may result from this kind of orientational interaction. This two-dimensional, agent-based model considers particles that move according to dynamical equations,

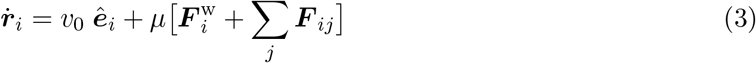

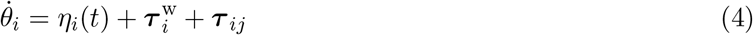

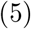

where position ***r***_*i*_ of the *i*^th^ particle with constant speed *v*_0_ along unit director 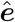. Particles experience repulsive-only harmonic interactions with a circular wall

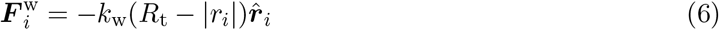

whenever *R*_t_ − |*r*_*i*_| < 0 and is elsewhere zero. Particles additionally experience repulsive-only interaction forces with other particles according to

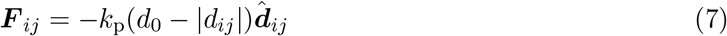

whenever *d*_0_ − |*d*_*ij*_| < 0 and is elsewhere zero.

Unit director 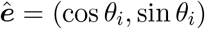 changes according to dynamical equation of its planar angle *θ*_*i*_ primarily according to uncorrelated Gaussian noise *η*_*i*_(*t*) defined with a mean of zero, ⟨*η*_*i*_(*t*)⟩ = 0 and variance *D*_*r*_, ⟨*η*_*i*_(*t*)*η*_j_(*t*′)⟩ = *D*_r_*δ*(*t* − *t′*). We model wall avoidance using a torque term

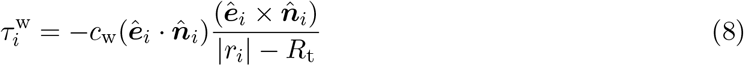

and is zero in the cases where 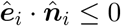 or (|*r*_*i*_| − *R*_t_) ≤ 0. In essentially the same way, we define avoidance interaction torques according to

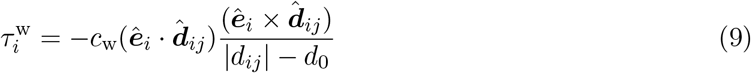

and is zero in the cases where 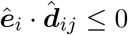 or (|*d*_*ij*_| − *d*_0_) ≤ 0.

Using this model, we simulate avoidance interactions at short range are able to turn them off by simply dropping the term *τ*_*ij*_.

#### 4.8.2 Simulations

We simulate our model using a standard Brownian dynamics algorithm.

## Supplementary Information

**Figure S1:**
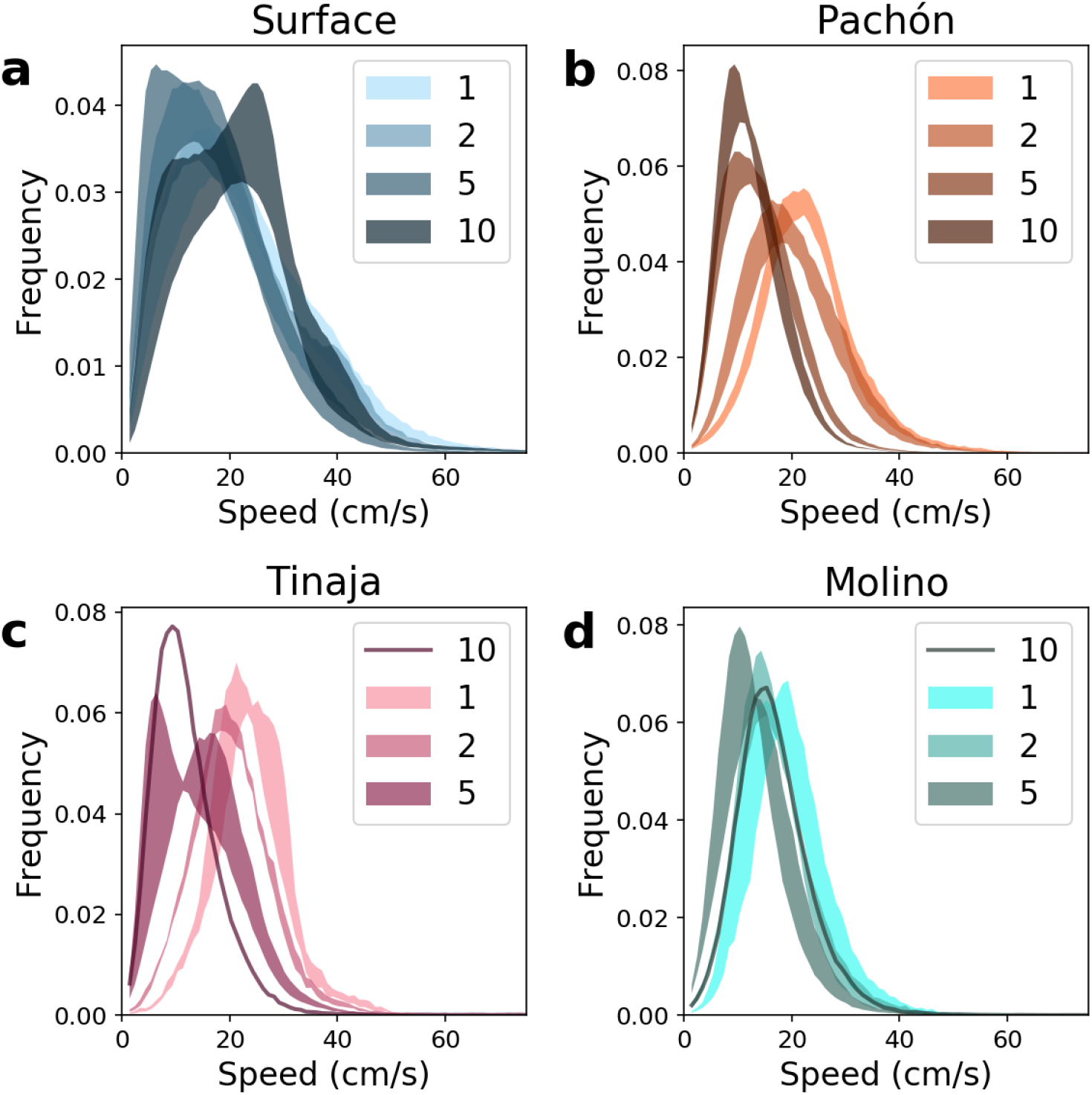
Speed distributions by group size for all fish populations, corresponding with Figure 3ab.

**Figure S2:**
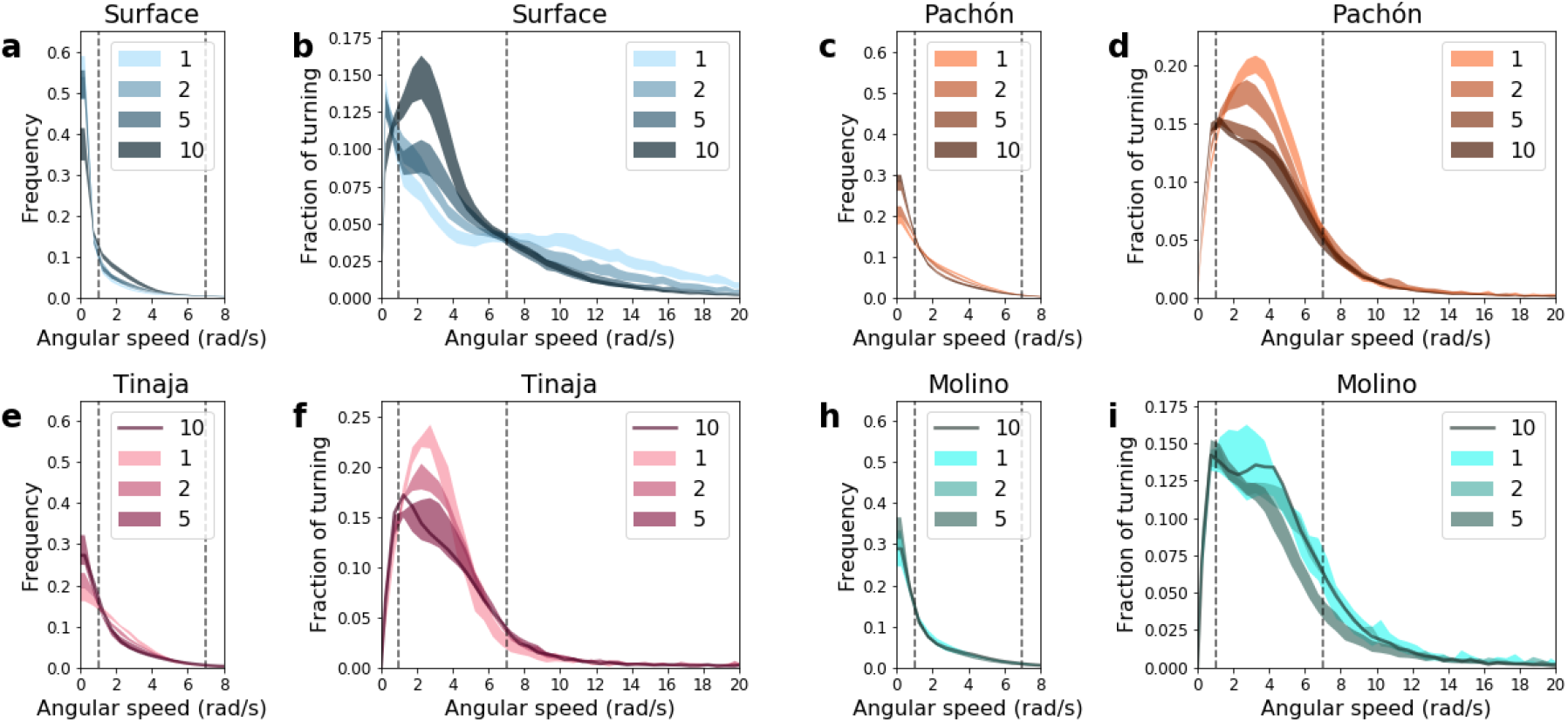
Turning distributions by group size for all fish populations, corresponding with Figure 4.

**Figure S3:**
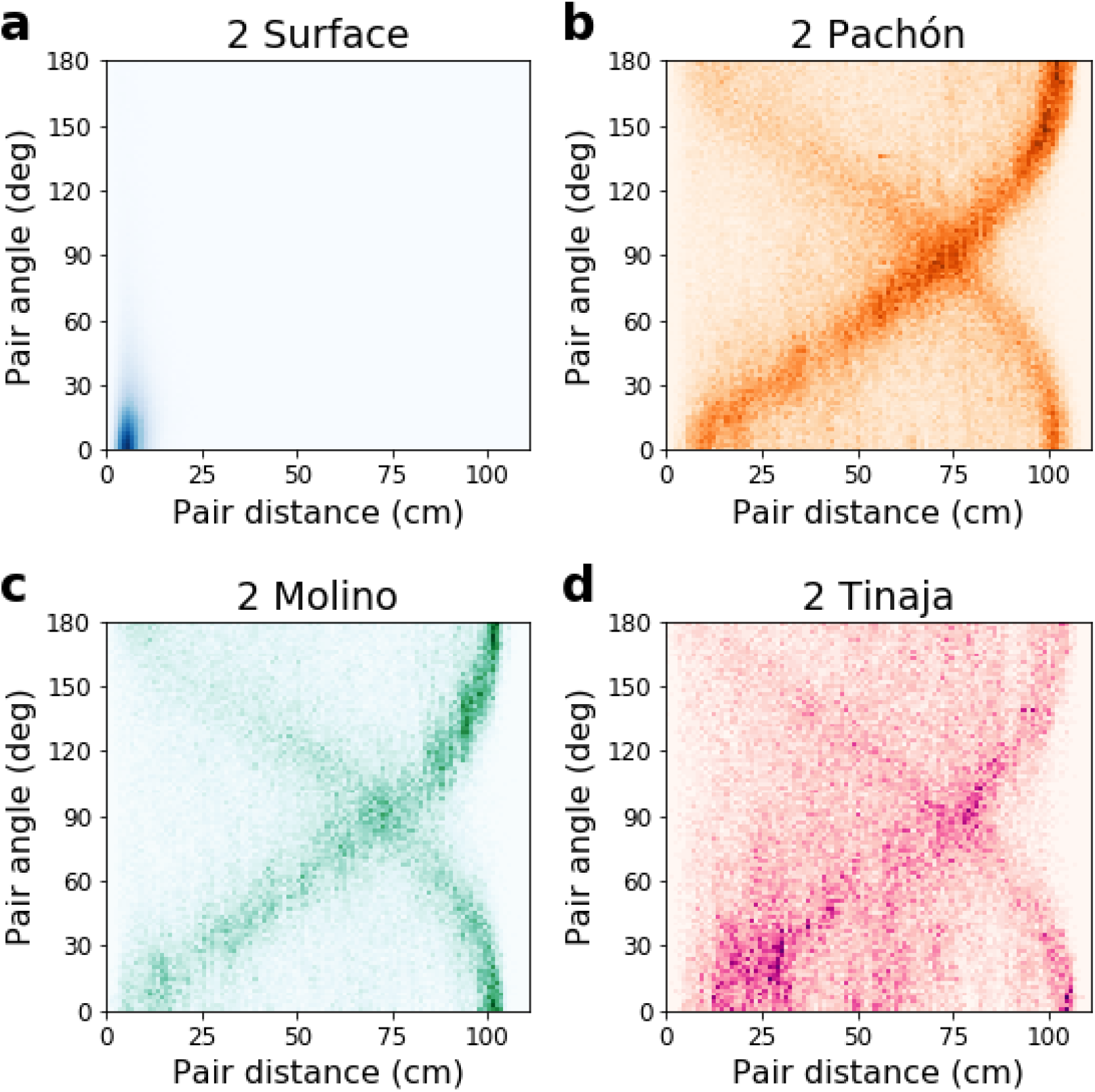
Pairwise maps of collective behavior for fish populations in groups of 2, corresponding with Figure 5bcef.

**Figure S4:**
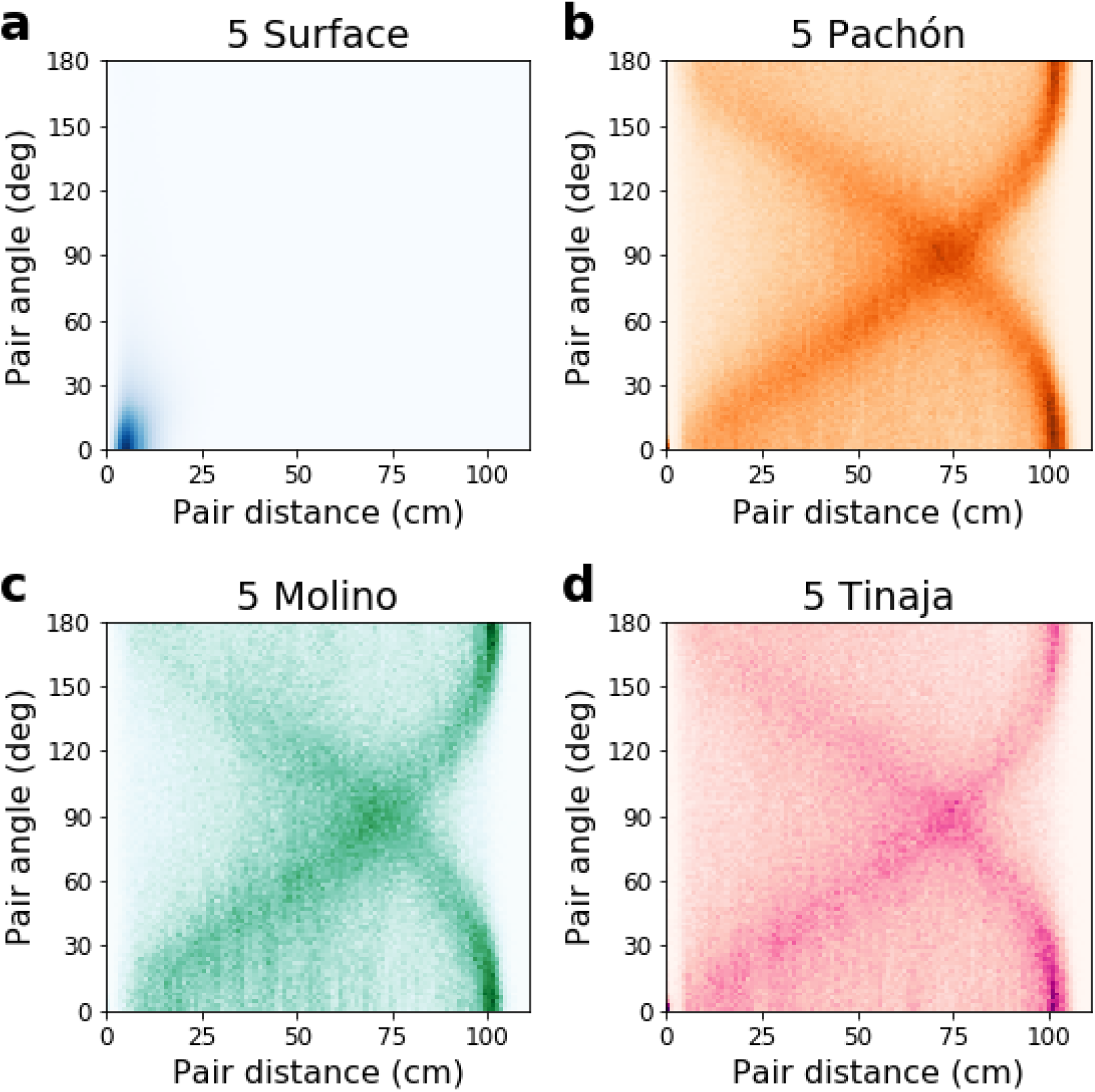
Pairwise maps of collective behavior for fish populations in groups of 5, corresponding with Figure 5bcef.

**Figure S5:**
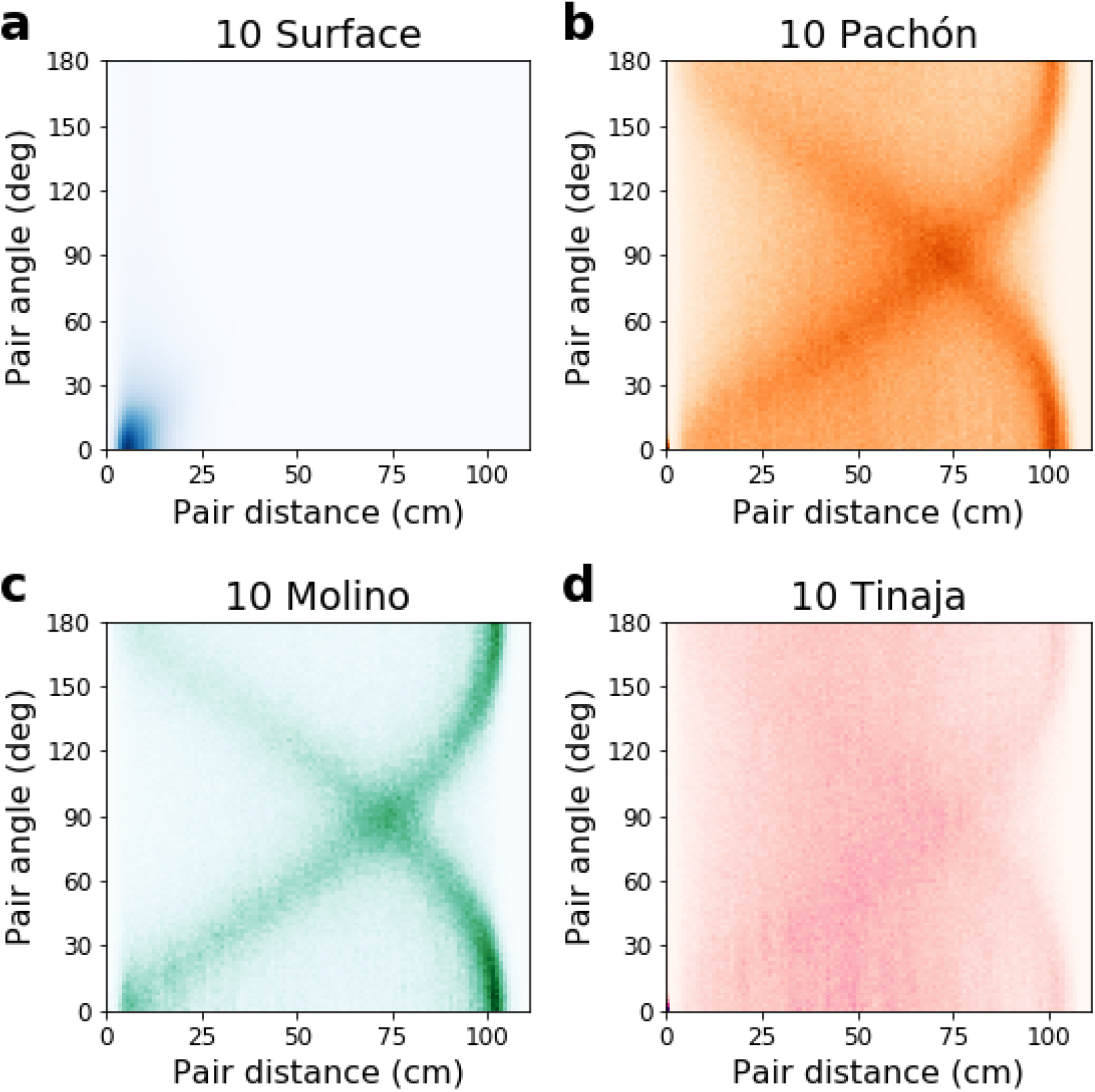
Pairwise maps of collective behavior for fish populations in groups of 10, corresponding with Figure 5bcef.

